# ApoE4 causes severe COVID-19 outcomes via downregulation of ACE2

**DOI:** 10.1101/2022.09.04.506474

**Authors:** Feng Chen, Yanting Chen, Qiongwei Ke, Yongxiang Wang, Xiongjin Chen, Xiaoping Peng, Yujie Cai, Shennan Li, Yuanhong Sun, Yao Ji, Yuling Jiang, Wenxian Wu, Yan Wang, Lili Cui

## Abstract

The coronavirus disease 2019 (COVID-19) pandemic is caused by severe acute respiratory syndrome coronavirus 2 (SARS-CoV-2); host cell entry by this virus relies on the interaction between the receptor-binding domain (RBD) of its spike glycoprotein and the angiotensin-converting enzyme 2 (ACE2) receptor on cell membranes. In addition to serving as a receptor for SARS-CoV-2, ACE2 was originally discovered as a protective factor in the renin–angiotensin system (RAS) that catalyses the degradation of angiotensin II (Ang II) to Ang 1-7, which is involved in multiple organ pathology. Recent genetic and clinical studies reported that ApoE4 expression is associated with increased susceptibility to SARS-CoV-2 infection and the development of severe COVID-19, but the underlying mechanism is currently unclear. In the present study, by using immunofluorescence staining, molecular dynamics simulations, proximity ligation assay (PLA) and coimmunoprecipitation (Co-IP) combined with a biolayer interferometry (BLI) assay, we found that ApoE interacts with both the spike protein and ACE2 but does not show obvious isoform-dependent binding effects. These data suggest that ApoE4 increases SARS-CoV-2 infectivity in a manner that may not depend on differential interactions with the spike protein or ACE2. Importantly, further immunoblotting and immunofluorescence staining results showed that ApoE4 significantly downregulates ACE2 protein expression *in vitro* and *in vivo* and subsequently decreases the conversion of Ang II to Ang 1-7, which could worsen tissue lesions; these findings provide a possible explain by which ApoE4 exacerbates COVID-19 disease.

## Introduction

The pandemic of coronavirus disease 2019 (COVID-19), which is caused by severe acute respiratory syndrome coronavirus 2 (SARS-CoV-2), has already resulted in more than 596 million confirmed cases and over 6 million related deaths worldwide as of August 26, 2022 (https://covid19.who.int). SARS-CoV-2 infection is initiated by the binding of the receptor-binding domain (RBD) of its spike protein to the angiotensin-converting enzyme 2 (ACE2) receptor(1), which is ubiquitously and widely expressed in lung, heart, brain, kidney, blood vessels, testis, the gastrointestinal tract, and other tissues(2). The ubiquitous expression of ACE2 renders these organs susceptible to SARS-CoV-2. In addition to being a major receptor for SARS-CoV-2, ACE2 was originally discovered to be a negative regulator of the renin–angiotensin system (RAS) that functions by catalysing the degradation of angiotensin II (Ang II) to Ang 1-7; thus, ACE2 is an important protective factor in multiple diseases, including hypertension, heart failure, myocardial infarction, acute lung injury, diabetes and Alzheimer’s disease (AD)(3-6). Over the course of the COVID-19 pandemic, emerging studies have reported that AD patients are particularly vulnerable to being infected by and spreading SARS-CoV-2, and once infected, these patients have significantly increased odds of mortality(7, 8). The COVID-19 incidence and case fatality rates differ among ethnicities, suggesting that genetic factors play an essential role in determining host responses to SARS-CoV-2(9). Recent genetic and clinical studies reported that patients in the UK Biobank Community Cohort who were homozygous for the *ApoE* ε4 gene had a 2.2-fold higher risk of developing severe COVID-19 independent of preexisting dementia, cardiovascular disease, and diabetes(10) as well as a 4.3-fold higher case fatality rate after COVID-19 than those who were homozygous for the *ApoE* ε3 gene(11). In Iranian group, a study observed that the ε4 allele increased the risk of the COVID-19 infection severity more than five times and the ε4/ε4 genotype showed a 17-fold higher in the risk of severe disease(12). *Shi* and colleagues reported that human-induced pluripotent stem cell (hiPSC)-derived ApoE4-expressing neurons and astrocytes are more susceptible to SARS-CoV-2 infection, and ApoE4 astrocytes exhibit a more severe response(13). The association between the ApoE4 genotype and the risk and severity of COVID-19 disease has also been reported by several other genetic studies(14-16), but the molecular mechanisms that are responsible for this isoform-dependent association remain poorly understood. Considering the strong effect of ApoE4 on COVID-19 severity and the high prevalence of the ε4 allele among individuals (the ε4 allele has a worldwide frequency of 13.7%), it is of great importance to explore the underlying mechanisms.

The human ApoE protein is composed of 299 amino acids, and it is abundantly expressed both in the central nervous system (CNS) and in the periphery, including the liver, brain, kidneys, adrenals, spleen, testis, skin, and lungs(17, 18); human ApoE functions as a primary regulator of cholesterol transport and lipid metabolism. Human ApoE has three common isoforms that differ by a single amino acid at residues 112 or 158; these isoforms are ApoE2 (Cys112 and Cys158), ApoE3 (Cys112 and Arg158), and ApoE4 (Arg112 and Arg158), and their differences obviously alter protein function(19). ApoE4 is considered to be the strongest genetic determinant for developing late-onset AD, and it is also known to be a risk factor for other CNS diseases, including Parkinson’s disease and Lewy body dementia(20, 21). In the periphery, ApoE4 was reported to be associated with an increased risk of <u>type 2 diabetes</u> and cardiovascular disease(22, 23). These ApoE4-related diseases have all been implicated in a higher risk of COVID-19(24, 25). In addition to SARS-CoV-2, ApoE gene polymorphisms are also associated with the attachment of and cellular and organismal responses to several other viruses, such as hepatitis C virus (HCV), HBV, and HIV-1(26-28). Notably, the role of ApoE in regulating viral infections seems to occur partly due to its interaction with heparan sulfate proteoglycans (HSPGs), functioning either as a Trojan horse or competing with viral particles for binding to HSPGs(29), suggesting that ApoE has the function of binding to the key receptor of virus and affecting virus infection.

In the present study, we aim to explore the underlying mechanism between ApoE isoforms and the severe COVID-19 risk. Here, we reported the potential interactions of different ApoE isoforms with the spike protein and ACE2, and further analysed the regulatory effects of different ApoE isoforms on the expression of ACE2, providing evidence that how ApoE4 increases the risk and severity of COVID-19.

## Results

### ApoE colocalizes with ACE2 *in vitro* and *in vivo*

Multiple lines of evidence indicate that SARS-CoV-2 enters human host cells via a high-affinity interaction with the ACE2 transmembrane receptor. SARS-CoV-2 has the remarkable ability to attack many different types of human host cells simultaneously. The lungs are the primary target of viral infection and replication, while kidney cells, neuronal cells and endothelial cells are also potential targets of SARS-CoV-2 infection, and the interaction of the spike protein with these cells may cause their dysregulation. Therefore, we first investigated whether ApoE and ACE2 were co-expressed in these tissues. The fluorescence results showed that ACE2 was primarily expressed on the cell membrane, and the colocalization between ApoE and ACE2 were observed in the lungs, kidneys, cortices and hearts of ApoE3-TR mice (Fig. 1A). Then the colocalization of ApoE and ACE2 were further observed in the A549, HEK-293T, SH-SY5Y, and HUVECs cell lines which corresponding to various tissues (Fig. 1B). The ApoE and ACE2 proteins were widely expressed in A549, HEK-293T, SH-SY5Y, and HUVECs cell lines, and ACE2 was expressed at the highest levels in HEK-293T cells (Fig. S1).

**Fig 1.**
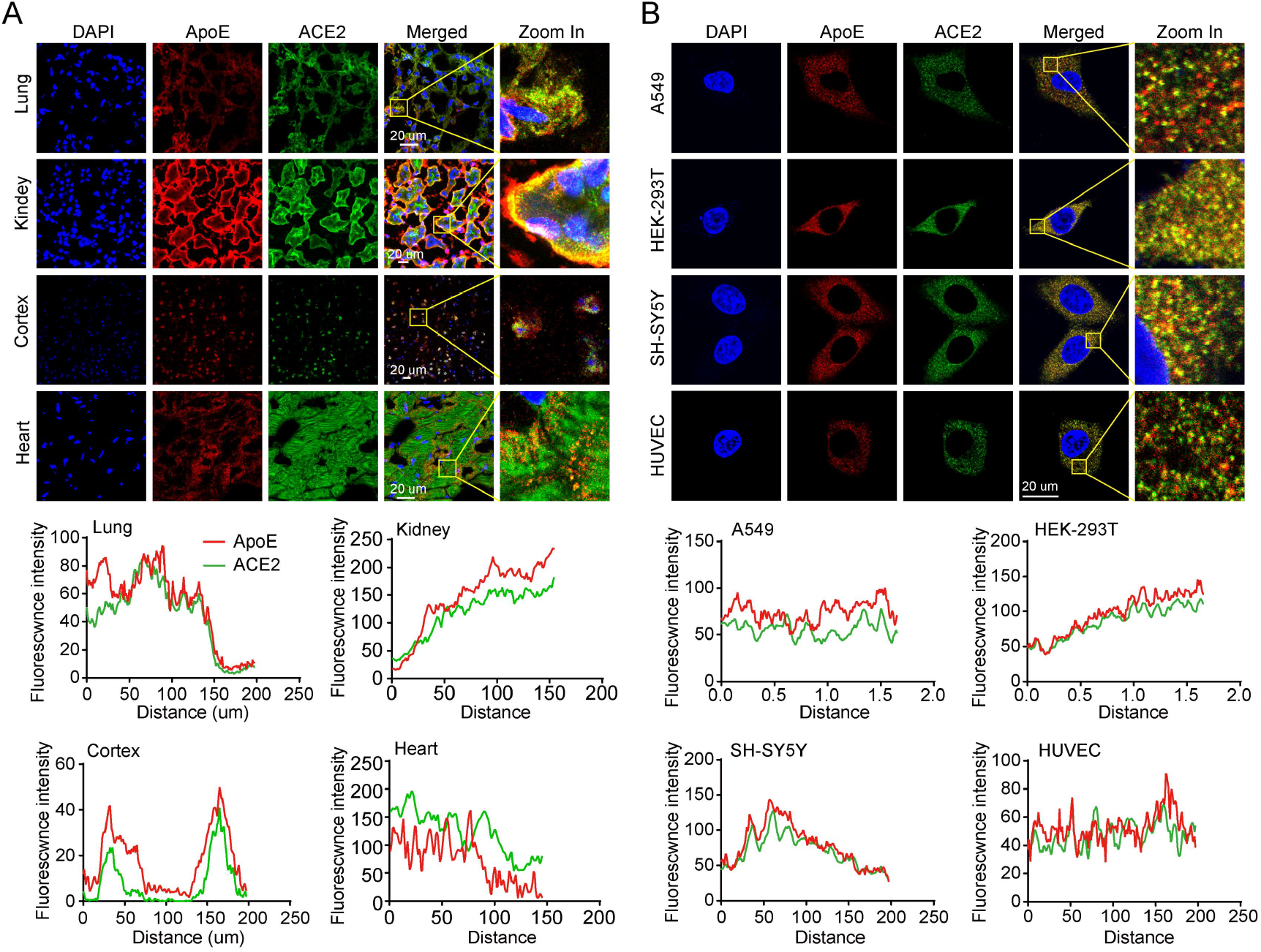
ApoE colocalizes with ACE2 *in vitro* and *in vivo*. (A) Representative coexpression of ApoE (red) and ACE2 (green) in lung, kidney, brain cortex and heart sections of ApoE3-TR mice as shown by immunofluorescence staining. The nucleus (blue) was stained with DAPI. Scale bars = 20 μm. (B) Representative coexpression of ApoE (red) and ACE2 (green) in A549, HEK-293, SH-SY5Y and HUVECs as shown by immunofluorescence staining. The nucleus (blue) was stained with DAPI. Scale bars = 20 μm.

### Molecular docking and simulation analysis of the interaction between ApoE and ACE2

To study the molecular interaction between the ApoE and ACE2 proteins, different ApoE isoforms were docked to the ACE2 protein by Rosetta software. The sites at which the ApoE proteins bound to the ACE2 protein were N53-K68, L45-K68 and N330-R357, and Y41-N64 and E329-D355 for ApoE2, ApoE3 and ApoE4, respectively. The binding positions of the ApoE isoforms with ACE2 protein overlapped with the region at which the spike protein interacts with ACE2 (Fig. 2A-C and Fig. S2A). Therefore, it can be hypothesized that the binding of ApoE to ACE2 may interfere with the binding of ACE2 to the spike protein, thereby affecting virus invasion. The RMSD represents the sum of all atomic deviations between the conformation at a certain moment and the target conformation and is an important basis for evaluating the stability of the system. As shown in Fig. S2B, the RMSD values of the ApoE proteins were stable after 40 ns of stimulation, with average values of 0.442 nm, 0.443 nm, and 0.475 nm for ApoE2, ApoE3 and ApoE4, respectively. The average RMSD of ACE2 in the three systems was 0.187 nm, 0.181 nm and 0.155 nm, respectively, with a relatively small fluctuation during the simulation process (Fig. S2C). Similar to RMSD, Rg indicates complex stability during the MD simulation. The average Rg values of the three systems during the simulation were 3.225 nm, 3.200 nm and 3.222 nm, respectively (Fig. S2D). Overall, the RMSD and Rg values of the three systems were not significantly different. We further assessed the changes in the solvent accessible surface area (SASA) of the three systems in the MD simulation process. After 20 ns of MD simulation, the SASA value of the ApoE2 system gradually stabilized, with an average value of 432 nm^2^, while the SASA values of the ApoE3 and ApoE4 systems decreased after 40 ns and stabilized after 80 ns, with average values of 417 nm^2^ and 418 nm^2^, respectively (Fig. S2E). In conclusion, there is no obvious difference in the SASA values among the three systems. Hydrogen bonding and hydrophobic interactions are the key forces that maintain the stability of interactions between biological macromolecules. The average numbers of hydrogen bonds between ACE2 and ApoE in the three systems were 4.47, 11.72 and 10.99, respectively (Fig. S2F), indicating that the hydrogen bonding between ApoE3 or ApoE4 and ACE2 is stronger than that between ApoE2 and ACE2. The mean values of the hydrophobic interaction between ApoE and ACE2 were 1.96, 8.45 and 3.67, respectively (Fig. S2G), suggesting that the hydrophobic interaction between ApoE3 and ACE2 is significantly stronger than those between ApoE2 or ApoE4 and ACE2. To further study the amino acid residue information of the interaction between the ACE2 and ApoE proteins, we analysed the protein binding mode after MD simulation. ApoE2 interacts with ACE2 with fewer amino acid residues and mainly relies on hydrogen bonding interactions (Fig. 2D). Specifically, N53, E56, Q60, N64, and K68 in ACE2 form hydrogen bonds with E179, R180, E244, Q248, and R251 in ApoE2, respectively. There is also a strong electrostatic interaction between E56--R180 and K68--E244. Therefore, it can be hypothesized that hydrogen bonding and electrostatic interactions are the main driving forces of the molecular interaction between ApoE2 and ACE2. Compared with the ApoE2-ACE2 system, the ApoE3-ACE2 system exhibited more hydrogen bonds between amino acid residues (Fig. 2E). E329, N330, N338, K353, G354, R357, N53, N58 and K68 in ACE2 form hydrogen bonds with E171, E179, L181, R191, R189 and E244 in ApoE3, respectively. In addition, there are some amino acid residues with strong hydrophobicity (L174, L181, L184, L45, T55, V339, etc.), which can form hydrophobic interactions and further enhance the affinity of the complex. Additionally, ApoE4 interacts with ACE2 at significantly more amino acid residues and with more hydrogen bonds than ApoE2 (Fig. 2F). Specifically, E35, E57, N58, N61, E329, N330, N338, Q340, and K353 in ACE2 form hydrogen bonds with E179, R178, R180, R189, R191, R240, and R251 in ApoE4, respectively. In addition, at the binding interface of the two proteins, there are some amino acid residues with strong hydrophobicity (P183, L184, M332, P336, V339, etc.), which can form hydrophobic interactions and further improve the affinity of the two proteins. Therefore, it can be hypothesized that hydrogen bonding and hydrophobic interactions are the main driving forces of these molecular interactions. The different binding modes may lead to the difference in binding energy. Therefore, we further analysed the change in the interaction energy between ACE2 and ApoE. The binding energies of the three systems fluctuated greatly in the early stage of the simulation, which occurred mainly because the complexes had not yet reached a stable state in the early stage of the simulation, and the interaction with the solvent was strong, so the intermolecular binding energy changed greatly. The binding energies of the three systems were basically stable after 80 ns, and the average values were −408 kJ/mol, −799 kJ/mol and −656 kJ/mol, respectively (Fig. S2H). The ApoE2-ACE2 system had a lower binding energy than the other systems, which could be the result of few hydrogen bonds and hydrophobic contacts.

**Fig 2.**
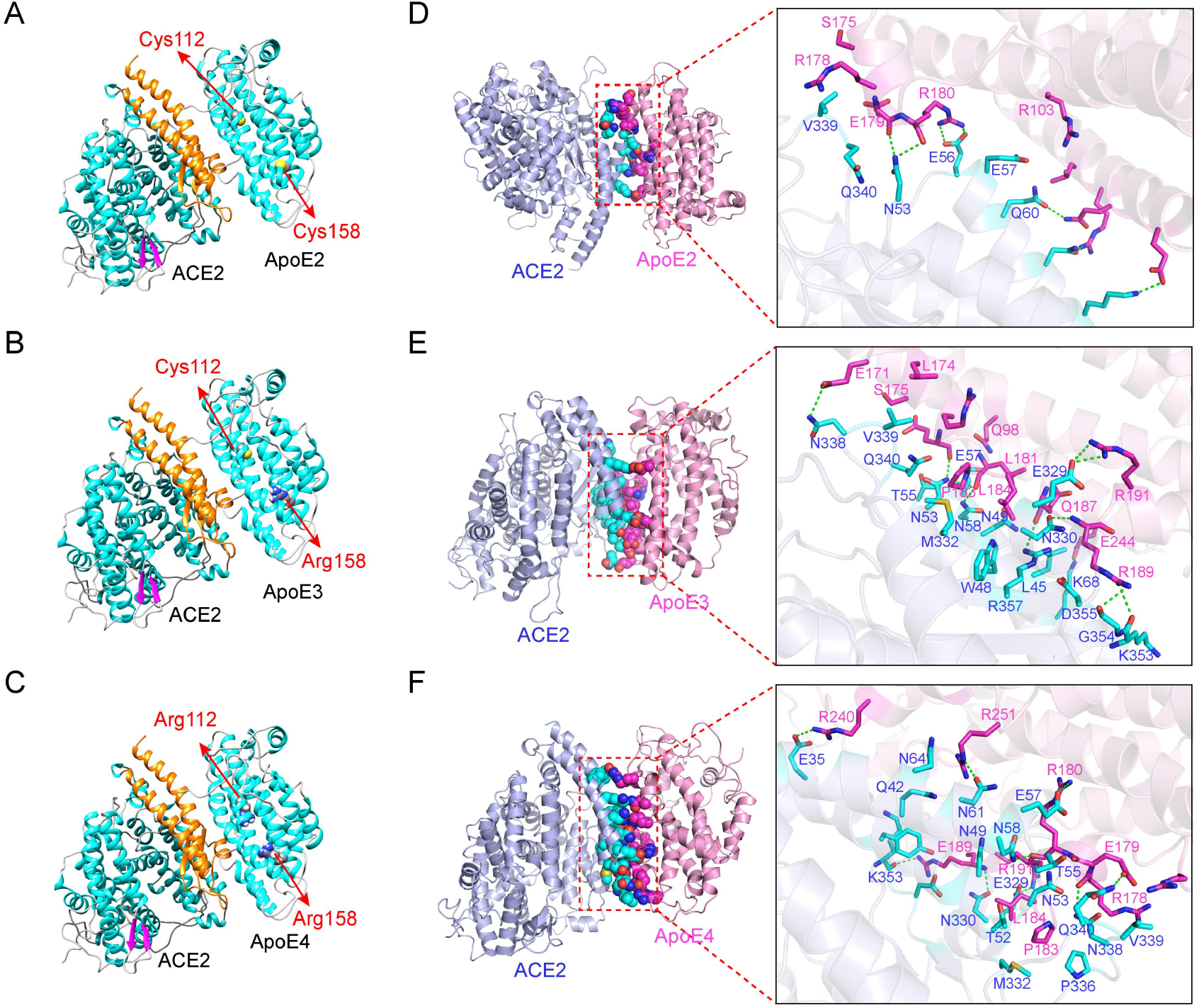
Molecular docking and simulation analyses of the interaction between ApoE and ACE2. (A-C) Predicted binding mode of ACE2 with ApoE2, ApoE3 or ApoE4 (the orange region of the ACE2 protein represents the region that binds to the S1 unit of the spike protein, and amino acids 112 and 158 of the ApoE protein are represented by spheres). (D-F) The three-dimensional mode and structural representation of the interface residues of the ApoE and ACE2 complexes (dotted green lines indicate hydrogen bonds, and blue and purple lines represent amino acid residues in ACE2 and ApoE proteins, respectively).

### ApoE interacts with ACE2 and the spike protein in an isoform-independent manner

Then, we assessed whether ApoE could interact with ACE2 in HEK-293T cells by PLA, which is a suitable method to describe protein-protein interactions at endogenous level with better quantitative precision. As shown in Fig. 3A, there were no significant different Duolink puncta (red) among ApoE isoforms, indicating a comparable binding between APOE isoforms and ACE2. The interaction was further evaluated by the Co-IP method in HEK-293T cells; in this experiment, Myc-tagged ACE2 was coexpressed with Flag-tagged ApoE (ApoE2, ApoE3 or ApoE4), and an anti-Myc antibody was used for immunoprecipitation, followed by western blotting analysis. The results showed that ApoE interacted with ACE2, while there was no significant difference between the ApoE isoforms (Fig. 3B). In an additional approach, streptavidin (SA) biosensors were used to precisely immobilize biotinylated ACE2, and then, its direct binding to purified recombinant ApoE was investigated (Fig. 3C). The BLI results showed that ApoE could directly bind to the ACE2 with a dose dependent manner, while there were no significant differently difference among ApoE isforms (Fig. 3D and Table S1). The spike protein of SARS-CoV-2 is responsible for binding to cellular receptors and subsequent viral entry into host cells. The spike protein encoded by the viral genome has two subunits, of which S1 contains the RBD that enables the virus to bind to its host target ACE2. Therefore, we further assessed the interaction between ApoE and the RBD of the spike protein, and the results showed that ApoE could bind to the spike protein with affinities of 1.71*10^−03^ mM for ApoE2, 5.89*10^−04^ mM for ApoE3, and 4.88*10^−04^ mM for ApoE4 (Fig. 3E and Table S2). Additionally, we observed dose- but not isoform-dependent binding. These results suggest that ApoE4 increases the risk of SARS-CoV-2 infection and that disease severity may not rely on the differential binding of ApoE isoforms to the spike protein and virus receptor ACE2.

**Fig 3.**
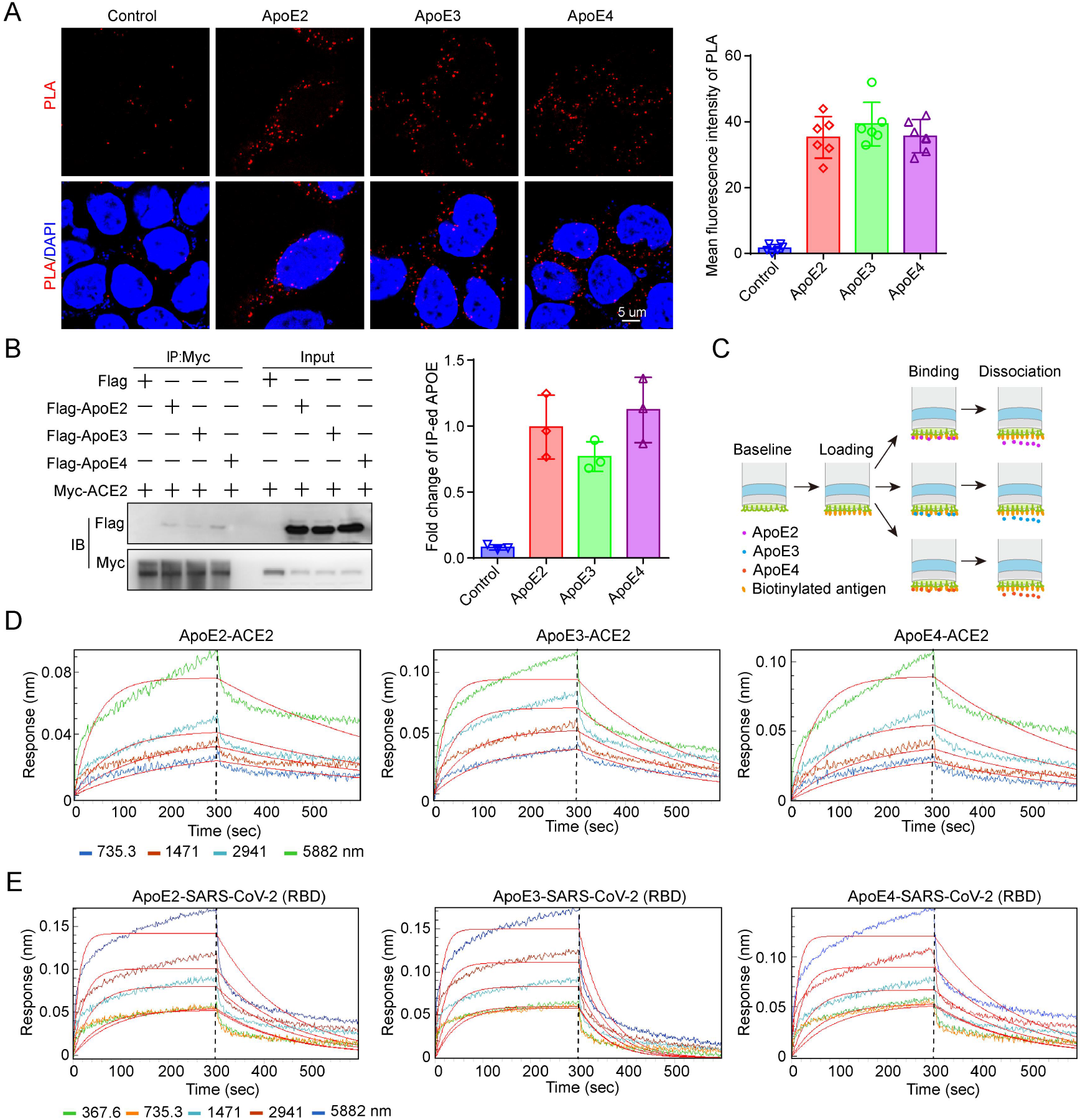
ApoE interacts with ACE2 and the spike protein in an isoform-independent manner. (A) Representative images of individual immunofluorescence staining of ApoE and ACE2 interaction tested by Duolink PLA in HEK-293T cells after 48 h of cotransfected by Myc-tagged ACE2 and 3×Flag-tagged ApoE. The red particles (ApoE/ACE2 interaction) represent their interaction. DAPI as a nuclear marker. Scale bar: 5 μm. (B) Plasmids carrying Myc-tagged ACE2 and 3×Flag-tagged ApoE (ApoE2, ApoE3 or ApoE4) were transiently cotransfected into HEK-293T cells. After 48 h of transfection, the cell lysates were immunoprecipitated with anti-Myc antibody and subsequently immunoblotted with anti-Myc and anti-Flag antibodies. The data are presented as the mean ± SD, *p < 0.05; **p < 0.01; ***p <0.001. (C) Schematic of the working principles of the BLI assay, which include loading, binding and dissociation. The binding affinities of ApoE and ACE2 were determined through BLI experiments. (D) The sensor surfaces were immersed in a solution of human ACE2 protein (20 µg/ml), and functionalized sensorgrams were captured upon incubation with human ApoE2, ApoE3, and ApoE4 at 735.3 (blue), 1471 (red), 2941 (light blue), and 5882 nM (green) (binding phase); then, the sensors were immersed in washing buffer (dissociation phase). (E) The binding affinities of ApoE and the SARS-CoV-2 RBD were determined through BLI experiments. The sensors were immersed in a solution of SARS-CoV-2 RBD (20 µg/ml), and functionalized sensorgrams were captured upon incubation with ApoE2, ApoE3, and ApoE4 at 367.6 (green), 735.3 (yellow), 1471 (light blue), 2941 (brown), and 5882 nM (blue); then, the sensors were immersed in washing buffer (dissociation phase).

### ApoE4 suppresses ACE2 expression *in vitro*

SARS-CoV-2 not only use ACE2 allowing virus entry but also downregulates ACE2 expression on cells, which may cause the imbalance between the RAS and ACE2/Mas axis and subsequently contribute to severe disease condition(30, 31). Consistently, studies reported a negative association between ACE2 amount and COVID-19 severity and fatality at both population and molecular levels(32). Thus, after analysing the interaction between ApoE and ACE2, we further evaluated the effect of ApoE gene polymorphisms on ACE2 expression at the cellular level, and the results showed that overexpression of ApoE4 led to a significant downregulation of ACE2 protein expression in SH-SY5Y, HEK-293T, A549 and HUVEC cells (Fig. 4A-D). Consistent results were obtained by immunofluorescence in these cells (Fig. 4 E-H). Considering the critical role of ACE2 in the RAS system, reduced expression of ACE2 may lead to dysregulation of RAS signalling, thereby contributing to disease progression.

**Fig 4.**
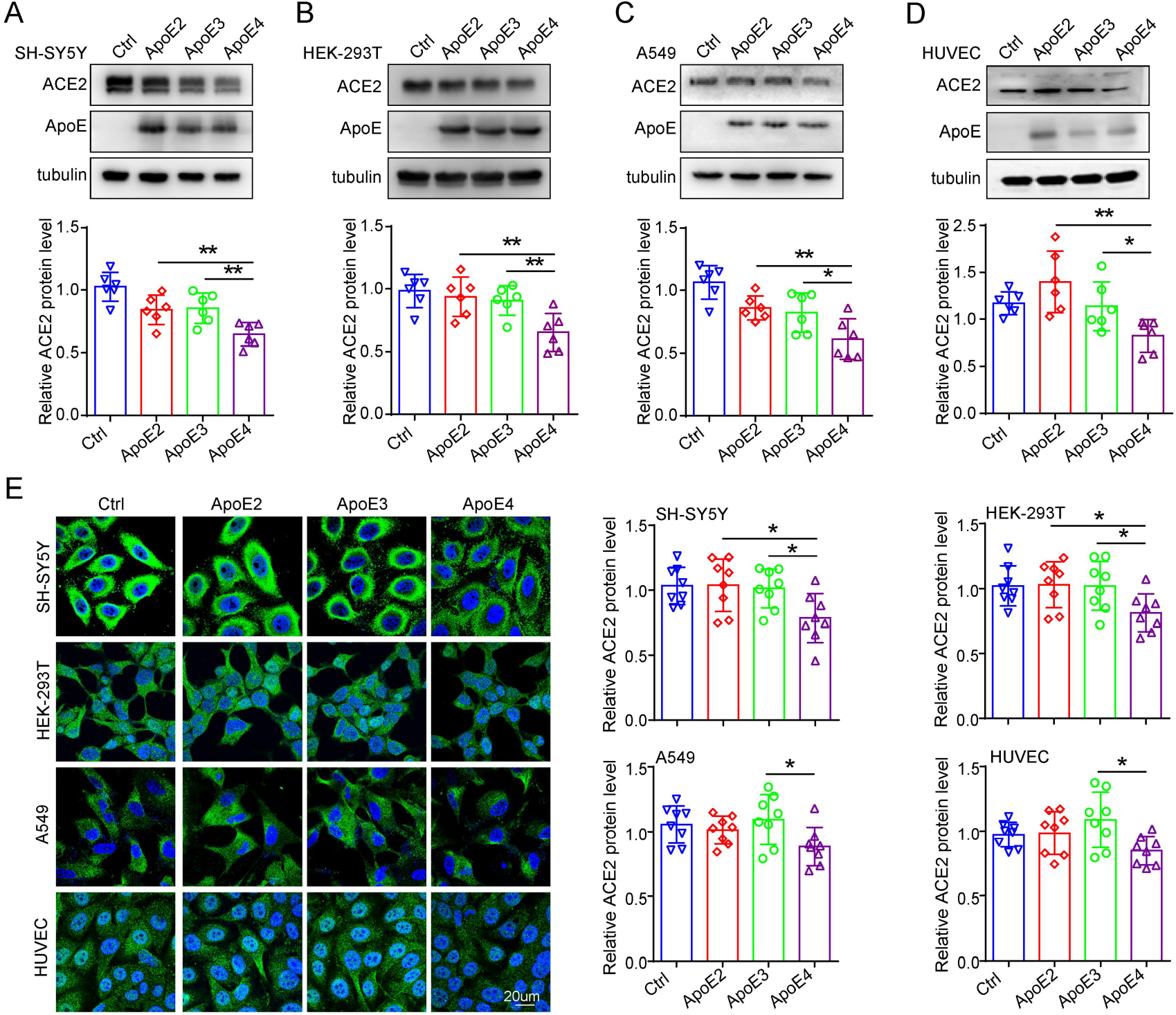
ApoE4 downregulates ACE2 protein expression *in vitro*. ACE2 protein levels in SH-SY5Y (A), HEK-293T (B), A549 (C) and HUVEC (D) cells were examined by western blotting after transfection with 1 µg/ml plasmids expressing Flag, ApoE2-Flag, ApoE3-Flag or ApoE4-Flag for 48 h. The results were normalized to the expression of a-tubulin. The data are expressed as the mean ± SD. (E-I) Cells were stained with ACE2 (green) antibody and counterstained with DAPI (blue) after transfection with 1 µg/ml Flag, ApoE2-Flag, ApoE3-Flag or ApoE4-Flag plasmids for 48 h. The data are expressed as the mean ± SD. Statistical differences were evaluated by one-way ANOVA. Scale bars, 20 μm. *p < 0.05; **p < 0.01; ***p <0.001.

### ApoE4 suppresses ACE2 expression *in vivo*

To determine whether this regulatory effect could be observed *in vivo*, here, we selected the ApoE-TR mouse model, in which expression of the human ApoE gene is regulated by the mouse ApoE promoter in mice on the C57BL/6J background. The mouse ApoE gene was replaced by the human ApoE2, ApoE3 or ApoE4 gene so that only the human ApoE gene is expressed. Thus, ApoE-TR mice are a good model for investigating the biological function of the human ApoE protein. First, we compared the protein expression of ACE2 in different tissues and found that ACE2 was abundantly expressed in bowel and kidney tissues (Fig. S3). Furthermore, the inhibition of ACE2 by ApoE4 was analysed in a variety of tissues, such as the cortex, hippocampus, liver, bowel, kidney, and heart. However, inhibitory effects were not observed in the lung tissues of ApoE-TR mice (Fig. 5). In addition, the regulatory relationship was further verified by immunofluorescence staining (Fig. S4). Considering that ACE2 serves as a crucial protective factor by catalysing the conversion of Ang II to Ang 1-7 in order to counter regulate the negative effects of RAS system, we further measured the Ang II and Ang 1-7 protein levels by ELISA. Compared with the ApoE2 and ApoE3 overexpression groups, overexpression of ApoE4 promoted the expression of Ang II and inhibited the expression of Ang 1-7 in HEK-293T cells (Fig. S5A and B). These results suggest that the ApoE polymorphism that is associated with increased COVID-19 severity may partly contribute to the effect of ApoE4-induced disorders in RAS signalling.

**Fig 5.**
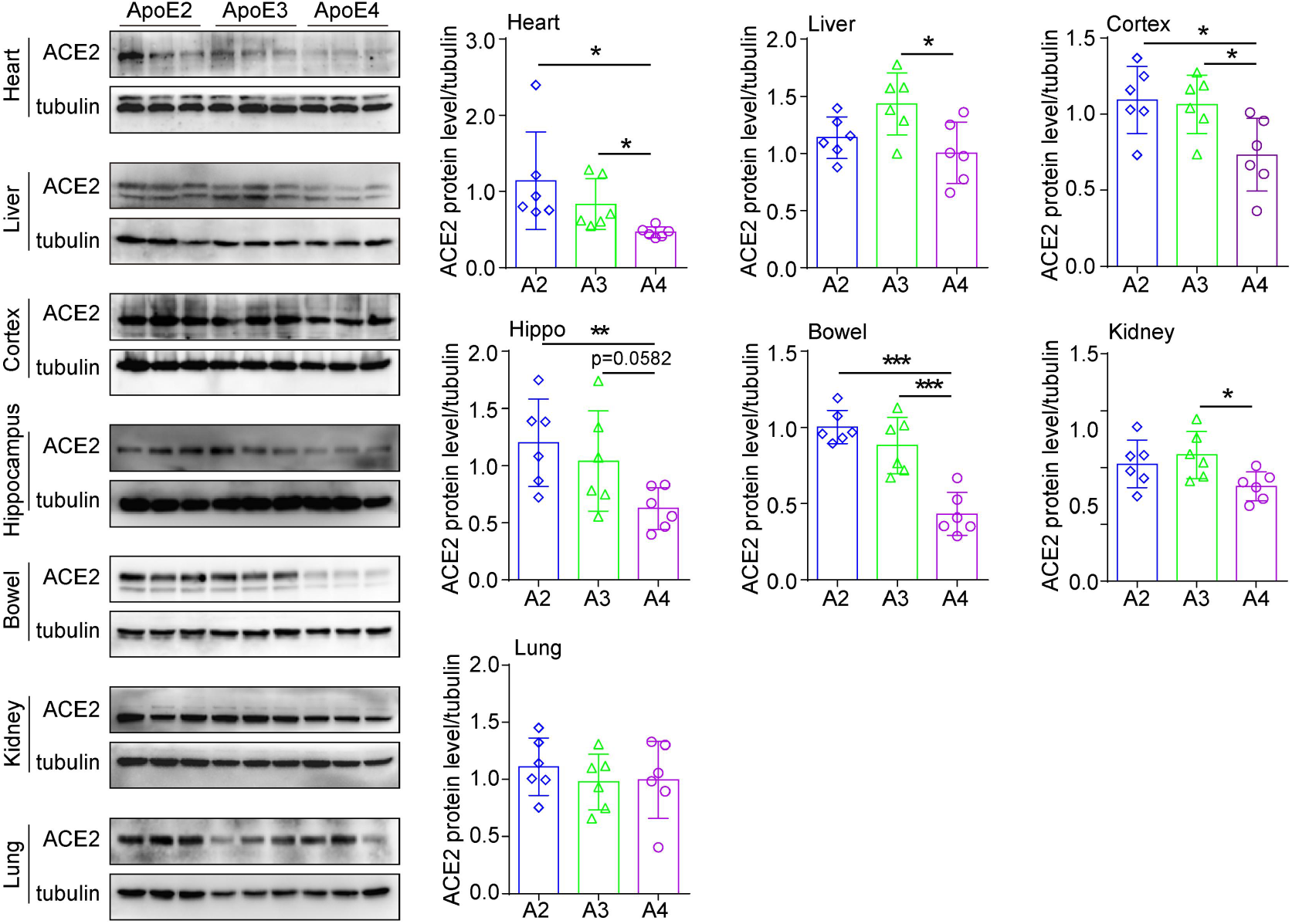
ApoE4 downregulates ACE2 protein expression *in vivo*. ACE2 protein levels in the cortex, hippocampus, liver, bowel, kidney, heart and lung of ApoE2-TR, ApoE3-TR and ApoE4-TR mice were measured by western blotting. The results were normalized to the expression of a-tubulin. n = 6 mice per group. The data are expressed as the mean ± SD. Statistical differences were evaluated by one-way ANOVA. *p < 0.05; **p < 0.01; ***p <0.001.

## Discussion

In the present study, we investigated the potential molecular link between ApoE and the spike protein and between ApoE and the SARS-CoV-2 primary receptor ACE2, and the results showed that ApoE interacts with both the spike protein and ACE2 but did not show isoform-dependent binding effects. Importantly, further results showed that ApoE4 significantly downregulates ACE2 protein expression *in vitro* and *in vivo* and consequently decreases the conversion of Ang II to Ang 1-7, which may provide a potential mechanism by which ApoE4 is associated with increased severity of COVID-19 (Fig. 6).

**Fig. 6.**
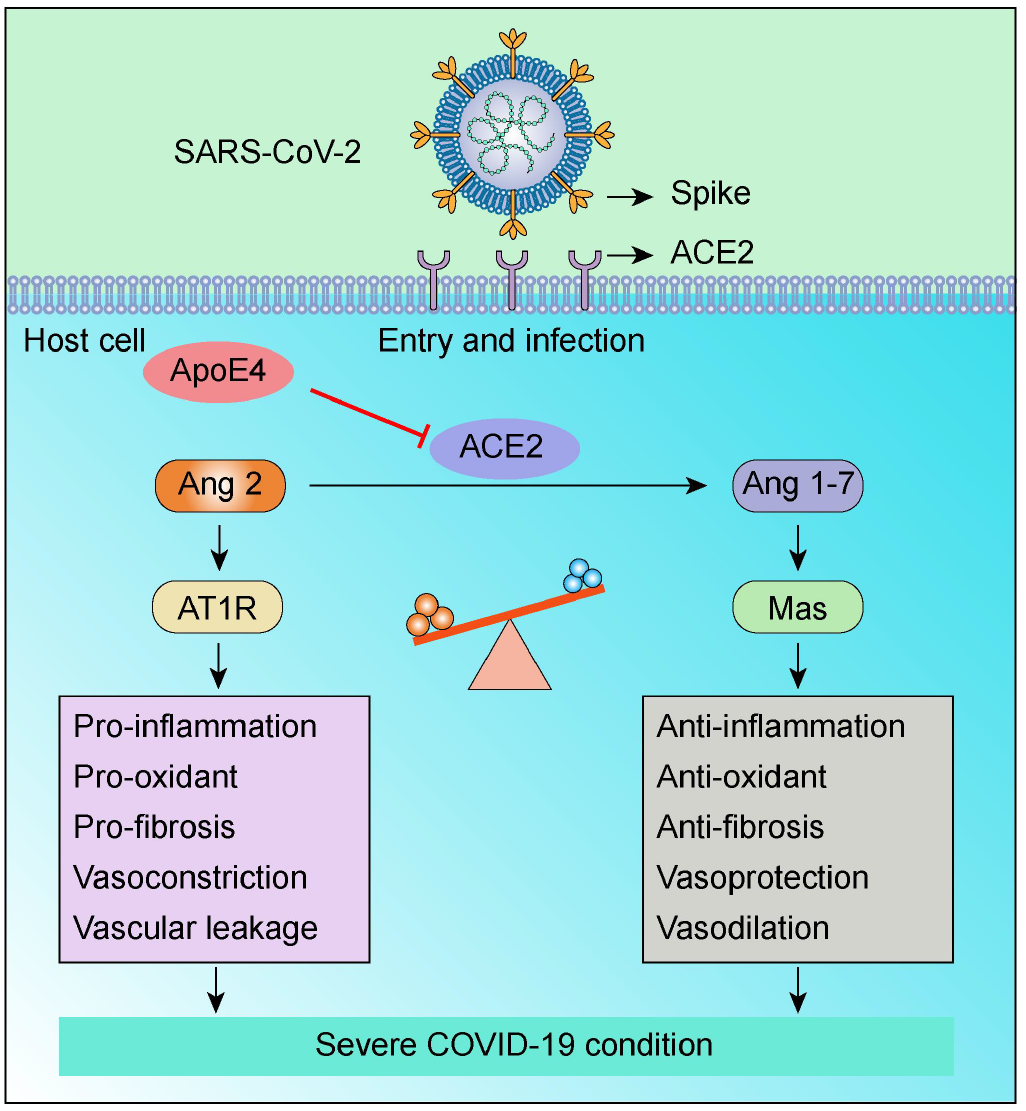
ApoE4 downregulates ACE2 protein expression and consequently decreases the conversion of Ang II to Ang 1-7, which may contribute to adverse COVID-19 outcomes.

SARS-CoV-2 entry into host cells is a crucial step in virus tropism, transmission, and pathogenesis. SARS-CoV-2 infection is initiated when the RBD of the spike protein binds to ACE2 receptors on cell membranes(33). Thus, neutralizing antibodies, ACE2-derived peptides, and small molecules that could bind to the spike protein or ACE2 have been investigated as promising therapeutic approaches for the treatment of COVID-19(34, 35), as these molecules disrupt the interaction between the spike protein and ACE2, which is the first step in viral infection. Emerging data have highlighted that genetic factors play an essential role in determining host responses to SARS-CoV-2. Recently, ApoE4 has been demonstrated to be associated with increased incidence and severity of COVID-19, but the mechanism remains elusive. Generally, ApoE primarily performs its biological functions by binding to lipids and receptors(36). Therefore, to investigate why ApoE4 increases the risk of SARS-CoV-2 infection and is associated with poor disease outcomes, it is important to investigate the potential interactions between ApoE and the spike protein and virus receptor. Although SARS-CoV-2 mainly attacks the respiratory system, organs such as the brain, kidneys, cardiovascular system, testes, liver, and intestine are all susceptible to SARS-CoV-2 infection, as ACE2 is broadly expressed in these organs(2, 37). Thus, we assessed the colocalization of ApoE and ACE2 in these cells and tissues. In addition, molecular docking results revealed favourable binding affinity between ApoE and ACE2, and the positions at which ApoE binds to ACE2 overlapped with the region at which the spike protein interacts with ACE2. Therefore, it can be hypothesized that the binding of ApoE to ACE2 may influence the binding of ACE2 to the spike protein, thereby affecting virus invasion. The direct interaction was further verified by Co-IP and BLI assays, and the binding affinities did not show an ApoE isoform-dependent manner. Our results are consistent with previous studies, in which the authors predicted that ApoE may interact with ACE2(38), and ApoE was coexpressed with ACE2 in type II alveolar cells(39), which exerts the most severe effect on virus pathology. Notable, recently Xu’s group also detected the association between ApoE and ACE2, and this interaction exhibits an inhibitory effect of SARS-CoV-2 pseudo-viral infection(16). Moreover, they found although ApoE3 and ApoE4 have a comparable binding affinity to ACE2, ApoE4 shows a lesser extent of pseudo-viruses cellular entry by spike docking onto the cell surface.

Previous studies have reported that ApoE can interact with virus proteins, such as proteins from HCV, HBV, and HIV. Can ApoE interact with SARS-CoV-2? In the present study, the interaction between ApoE and the RBD of the spike protein was explored by BLI assay. The data showed that ApoE directly bound to the spike protein in a dose-dependent manner, while there were also no obvious differences among the ApoE isoforms, suggesting that ApoE4 increases the risk of SARS-CoV-2 infection in a manner that may not rely on differential binding between ApoE isoforms and the spike protein. The results were consistent with and an extension of a recent study in which molecular docking and molecular dynamics simulation data suggested that there may be an interaction between ApoE and the spike protein and that such an interaction may cause structural changes in ApoE, which may activate the ApoE metabolic pathway and facilitate SARS-CoV-2 entry(40).

ACE2 is a type I transmembrane glycoprotein that is widely expressed throughout the human body(41). ACE2 is much more than just the primary receptor for SARS-CoV-2, and it was originally discovered as a protective factor involved in the RAS pathway. The predominant function of ACE2 is to metabolize Ang II to Ang 1-7, reduce inflammation and oxidative damage, and counteract the harmful effects of the ACE-Ang II-AT1 pathway; therefore, ACE2 performs multiple salutary biological functions in several diseases, such as heart failure, myocardial infarction, hypertension, kidney diseases, acute lung injury, diabetes and AD(3-6). Thus, we further examined the effect of different ApoE genotypes on ACE2 protein expression. Immunoblotting and immunofluorescence staining results showed that ApoE4 significantly downregulated ACE2 protein expression at the cellular level and in the organs of ApoE-TR mice. Consistent results from an autopsy study reported that ACE2 activity was decreased by almost 50% in the brain tissues of AD patients compared with age-matched controls, and the reduction was associated with the presence of an ε4 allele(42). Specifically, ACE2 activity was inversely associated with the levels of Aβ and phosphorylated tau pathology. As an indirect measurement of reduced ACE2 activity, the amount of Ang II was elevated in the cells that overexpressed ApoE4. ACE2 reduces adverse effects by degrading Ang II peptides, thereby eliminating or diminishing their deleterious potential, and by generating Ang 1-7, which performs a variety of beneficial functions that oppose the functions of Ang II through efficient interaction with the G protein coupled receptor Mas and Ang II type 2 (AT2) receptors(43). Thus, we provide a possible mechanism by which ApoE4 increases the severity of COVID-19 infection.

It seems paradoxical that the ApoE4-mediated decrease in ACE2 levels would decrease SARS-CoV-2 infectivity, thereby exerting a protective effect. In theory, ACE2 downregulation might reduce the risk of SARS-CoV-2 infection due to decreased availability of the virus receptor. However, studies have suggested that ACE2 deficiency can exacerbate tissue lesions because of the major imbalance in the RAS(44). It is important to note that SARS-CoV-2 uses ACE2 to enter cells and downregulates ACE2 expression, which in turn results in the loss of the beneficial effects of the ACE2/Mas axis in the lungs, cardiovascular system, kidney, and other tissues, leading to adverse COVID-19 outcomes(30, 31). Therefore, reduced ACE2 protein expression is associated with a poor prognosis in patients with SARS-CoV-2(32, 45). Clinical observational studies have shown that in most cases, respiratory distress occurs many days after infection, indicating that this may not be a direct effect of the initial viral infection but rather an effect of the host response to the loss of ACE2 function and dysregulation of the Ang II/ACE2 pathways(46). In addition, it is interesting to note that several conditions associated with viral infection and disease severity share a variable degree of ACE2 deficiency. Given the above premises, it is tempting to hypothesize that ApoE4 inhibits the protein expression of ACE2, resulting in ACE2 downregulation, which reverses its protective effect and may lead to unfavourable outcomes in patients with COVID-19.

Several limitations in our study should be mentioned. First, although there were no obvious differences in the binding abilities of the ApoE subtypes to the spike and ACE2 proteins, the total amount of bound ApoE protein may be different *in vivo*, since ApoE protein levels have been reported to be reduced by the inheritance of the ε4 allele(47, 48), and this difference may contribute to the predisposition to the risk and prognosis of SARS-CoV-2 infection. In addition, whether the binding of the ApoE subtypes to the spike and ACE2 proteins interferes with the interaction between the spike and ACE2 proteins should be determined. Since ApoE can interact with both ACE2 and the spike protein, we could not determine the effect of ApoE on the ability of ACE2 and the spike protein to bind based on BLI or competitive ELISA analyses. Therefore, it cannot be completely ruled out that ApoE polymorphisms affect the risk of viral infection by interfering with the interaction of the spike protein with the ACE2 receptor. Whether different ApoE subtype affect the level of ACE2 by affecting the binding with ACE2, this causal relationship is still possible and needs further exploration. Moreover, in the present study, we were unable to establish a causal relationship between ApoE4-induced ACE2 downregulation and COVID-19 severity. ApoE4 may exert multifaceted effects in patients with COVID-19, such as increased blood–brain barrier permeability(49), promotion of inflammatory responses(50), and decreased expression of several antiviral defence genes(51). These factors have also been reported to be associated with poor clinical outcomes in patients with COVID-19. Therefore, future studies are warranted to determine whether ApoE4 increases the risk of COVID-19 progression independent of these factors.

Collectively, we provided a potential mechanism by which ApoE4 increases the severity of COVID-19 infection. Our results showed that ApoE can interact with both the spike protein and ACE2 in an isoform-independent manner. And ApoE4 can significantly inhibit ACE2 expression level compare with the other ApoE isoform in vivo and in vitro, that may explain why the APOE4 carrier have a poor prognosis in patients with SARS-CoV-2. However, future studies are needed to detect how ApoE gene polymorphism affects ACE2 protein expression and other mechanisms by which ApoE gene polymorphism affects SARS-CoV-2 infection and disease severity.

## Materials and Methods

A detailed description of materials and methods is provided in Supplemental information

## Data Availability

All study data are included in the article and/or supporting information.

## Acknowledgments

This work was supported by the National Natural Science Foundation of China (81671181, 82101269), Guangdong Basic and Applied Basic Research Foundation, (2021A1515110965) and the Project funded by China Postdoctoral Science Foundation (2022M710848).

## Author contributions

LC and YW designed research; FC, YTC, QK, YW, XC, YJC, SL, YS, YJ, YLJ, XP and WW performed research; FC, YTC and QK analyzed data; LC and FC wrote the paper.

## Supplemental information

### Materials and methods

#### Animals

Human ApoE-targeted replacement (ApoE-TR) mice, in which the expression of human ApoE is controlled by the mouse ApoE promoter, where generated on the C57BL/6J background. The mouse ApoE gene was replaced by the human ApoE2, ApoE3 or ApoE4 gene so that only the human ApoE gene is expressed. ApoE-TR mice were provided by Cyagen Biosciences and housed in the Laboratory Animal Center (SPF grade) of Guangdong Medical University. All the mice were maintained under a 12 h–12 h light/dark cycle and had free access to food and water. All the animal experimental procedures were performed in accordance with the Guide for the Care and Use of Laboratory Animals and approved by the laboratory animal ethical committee of Guangdong Medical University.

#### Cell culture and transfection

Human embryonic kidney (HEK)-293T cells, human neuroblastoma cells (SH-SY5Y), human lung cancer cells (A549) and human umbilical vein endothelial cells (HUVEC) were cultured in Dulbecco’s modified Eagle’s medium (DMEM) supplemented with 10% foetal bovine serum (FBS), 100 U/ml penicillin and 100 mg/ml streptomycin. The cells were maintained at 37°C with 5% CO_2_. pCMVPuro05-Control, pCMVPuro05-ApoE2, pCMVPuro05-ApoE3 and pCMVPuro05-ApoE4 plasmids (1 µg/ml) were transfected into these cells (at ∼70% confluence) using a Lipofectamine 3000 transfection kit (Invitrogen). Six hours after transfection, the cell medium was replaced with DMEM. After 48 h, the cells were harvested for western blotting analysis or immunofluorescence staining.

#### Western blotting analysis

Total cell and tissue proteins were extracted with RIPA buffer (Solarbio, R0010) supplemented with Protease Inhibitor Cocktail Tablets (Roche) and PMSF (Boster), and the protein concentrations were quantified by a Pierce™ BCA Protein Assay Kit (Thermo, TL276863). Equal amounts of proteins were separated by SDS–PAGE and transferred to PVDF membranes (A10122278, GE, USA). After blocking with 5% nonfat milk and washing with Tris-buffered saline with Tween 20 (TBST) buffer, the membranes were incubated with anti-ApoE (1:1,000, Meridian, K74180B) or anti-ACE2 (1:1,000, Abcam, ab15348) antibodies at 4°C overnight. Then, the membranes were washed with TBST and incubated with secondary antibodies at room temperature for 1 h. Finally, the immunoreactive bands were visualized by enhanced chemiluminescence (Invigentech). Protein expression was quantified by the measuring the band densities using ImageJ software. α-Tubulin (Abcam, ab4074) was used as the loading control.

#### Immunofluorescence staining

Tissue sections and cell samples were fixed with 4% paraformaldehyde solution for 15 min, washed with PBS, incubated with permeabilization agent for 5 min, blocked with 10% goat serum for 30 min at room temperature, and then incubated with anti-ACE2 antibody (1:500, Abcam, ab15348) and anti-ApoE (1:300, Santa Cruz, sc-390925) antibodies overnight at 4°C. The next day, the cells were washed with PBS and incubated with fluorescent secondary antibodies (Alexa Fluor® 488, 1:400, Abcam; Alexa Fluor® 647, 1:400, Abcam) for 1 h at room temperature. The nuclei were stained with 4’,6-diamidino-2-phenylindole (DAPI) for 3 min. Fluorescence signals were acquired by confocal microscopy (FV3000, Olympus).

#### Proximity ligation assay

PLA was performed using the Duolink *in situ* Red Starter Kit Mouse/Rabbit (Sigma-Aldrich, F1635) following the manufacturer’s instructions. HEK-293T cells were cotransfected with 1 µg/ml pCMV-ACE2-MYC and pCMVPuro05-Control, pCMVPuro05-ApoE2-3×Flag, pCMVPuro05-ApoE3-3×Flag or pCMVPuro05-ApoE4-3×Flag. After transfection for 48 h, the cells were incubated with anti-Flag (Abbkine, A02010) and anti-ACE2 (Abcam, ab15348) primary antibodies overnight at 4°C. The cells were then incubated with anti-mouse MINUS and anti-rabbit PLUS proximity probes for 1 h at 37°C. Ligation and amplification were performed using the Duolink *in situ* detection reagent kit according to the manufacturer’s protocol. Finally, the cells were mounted on a slide with the Duolink *in situ* mounting medium with DAPI. Images were captured using an Olympus FV1000 confocal microscope, and the red spots represent the interactions between ApoE and ACE2.

#### Coimmunoprecipitation (Co-IP) assay

For the Co-IP analysis, HEK-293T cells were cotransfected with 1 µg/ml pCMV-ACE2-MYC and pCMVPuro05-Control, pCMVPuro05-ApoE2-3×Flag, pCMVPuro05-ApoE3-3×Flag or pCMVPuro05-ApoE4-3×Flag. After transfection for 48 h, the cells were washed with PBS and lysed with IP lysis buffer for 30 min at 4°C. The supernatants of the cell lysates were incubated with the corresponding antibody overnight at 4°C, incubated with protein A/G agarose beads (Santa Cruz, sc-2003) for 2 h at 4°C, and washed three times with lysis buffer. The proteins were analysed by western blotting using anti-Flag (1:1,000, Abbkine, A02011) and anti-MYC (1:1,000, Beyotime, AM926) antibodies.

#### Biolayer interferometry (BLI) binding assay

BLI binding experiments were performed using Octet RED96 equipment (ForteBio). The recombinant RBD (SinoBiological, 40592-V05H) and recombinant human ACE2 proteins (SinoBiological, 10108-H02H) were captured with an anti-mIgG Fc Capture (AMC) probe in PBST solution and then incubated with recombinant human ApoE2 (PeproTech, 350-12-500UG), ApoE3 (PeproTech, 350-02-500UG) and ApoE4 (PeproTech, 350-04-500UG) proteins. Then, the fully reacted solid-phase conjugates were dissociated in PBST buffer for analysis. The kinetic values were fitted to a 1:1 Langmuir binding model. The results were analysed with ForteBio Data Analysis 11.0 software to determine the association rate, dissociation rate and affinity constant.

#### Molecular docking and molecular dynamics simulation

The crystal structure of human ApoE3 (PDB ID: 2L7B) was obtained from the Protein Data Bank (PDB). The three ApoE isoforms differ from one another only at positions 112 and 158 (ApoE2: Cys112 and Cys158; ApoE3: Cys112 and Arg158; ApoE4: Arg112 and Arg158). Therefore, the Arg residue at 158 was mutated to Cys, and the Cys residue at 112 was mutated to Arg to obtain the initial structures of the ApoE2 and ApoE4 proteins in PyMOL 2.1. The structure of the ACE2 protein was obtained from PDB (PDB ID: 6M0J). The Rosetta tool was used to dock ACE2 to ApoE2, ApoE3 or ApoE4, and 1000 conformations were collected. Finally, a reasonable docking structure was selected. Unreasonable atomic contacts were released by using energy-optimized methods. The Amber14sb force field was used for energy optimization. First, the 2000-step steepest descent method was used to optimize the structure, and then, the 2000-step conjugate gradient method was used to further optimize the structure. The final results were used for subsequent analysis. The GROMACS 2019.6 program was used for molecular dynamics simulation under constant temperature and pressure and periodic boundary conditions. During the molecular dynamics simulation, hydrogen bonds were constrained using the LINCS algorithm with an integration step size of 2 fs. Electrostatic interactions were calculated using the Particle–mesh Ewald (PME) method. The nonbonded interaction cut-off was set to 10 Å and updated every 10 steps. The V-rescale temperature coupling method was used to maintain the simulated temperature at 298 K, and the Parrinello-Rahman method was used to maintain the pressure at 1 bar. The hydrogen bond adopted the geometric criterion, the angle between the hydrogen bond donor and the acceptor was greater than 130°, and the distance was less than 0.35 nm. Energy minimization was performed using the steepest descent method to eliminate excessively close contacts between atoms; then, a 100 ps NPT equilibrium simulation was performed; finally, a 100 ns molecular dynamics simulation was performed for each of the three systems, and conformations were saved every 50 ps. The simulation results were visualized using the Gromacs embedded program and VMD.

#### Enzyme-linked immunosorbent assay (ELISA)

After 48 h of transfection, the culture media were collected and centrifuged for 10 min at 4°C at 3,000 rpm. The protein levels of Ang II (Cusabio, CSB-E04500h) and Ang 1-7 (Cusabio, CSB-E14242h) were measured with ELISA kits according to the manufacturer’s guidelines. The OD value at 450 nm was measured with a microplate reader (Bioteck, USA). The minimum detectable levels of human Ang II and Ang 1-7 were 9.75 pg/ml and 1.95 pg/ml, respectively.

#### Statistical analysis

The data are presented as the mean ± standard deviation (SD). Statistical significance between two groups was determined by unpaired two-tailed Student’s t test. One-way ANOVA was used to analyse the differences in a single independent variable between two groups. Multiple group comparisons were performed using two-way ANOVA. All western blotting and immunofluorescence data were obtained from at least three replicates. All the data and graphs in this paper were analysed and generated using GraphPad Prism 9.0. A value of P<0.05 was considered statistically significant (*P<0.05, **P<0.01, ***P<0.001).

## Supplemental Figures

**Fig S1.**
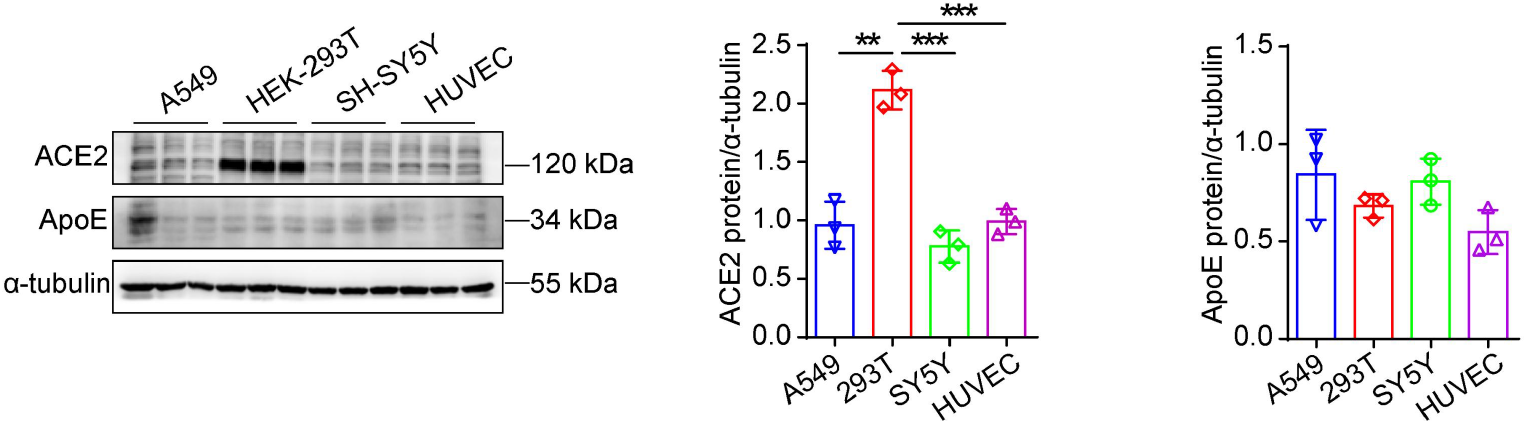
Expression of ApoE and ACE2 *in vitro*. Representative western blotting analysis of the ApoE and ACE2 protein levels in A549, HEK-293T, SH-SY5Y, and HUVECs; the data are shown as the mean ± SD of three independent experiments. α-Tubulin was used as a loading control. *P* values were calculated using one-way ANOVA, \**p* < 0.05; \*\**p* < 0.01; \*\*\**p* <0.001.

**Fig S2.**
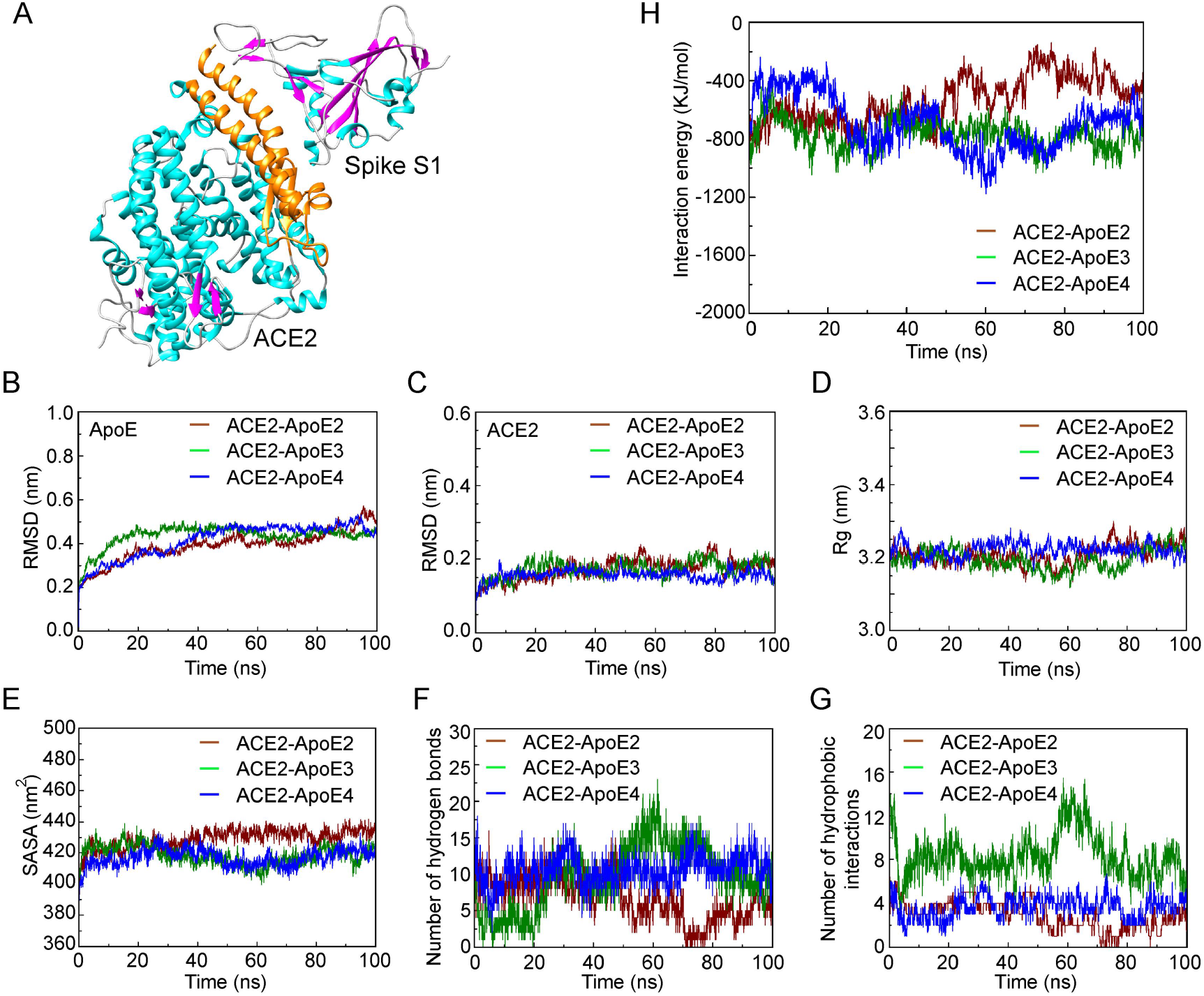
Molecular docking and simulation analyses of the interaction between ApoE and ACE2. (A) Molecular docking simulation of the SARS-CoV-2 RBD interacting with ACE2 (the orange region of the ACE2 protein represents the region that binds to the spike S1 protein). (B and C) Plot of backbone RMSD versus time (ns) for ApoE and ACE2. (D) Plot of Rg versus time (ns) for ApoE-ACE2. (E) Plot of total SASA versus time (ns) for ApoE-ACE2 complexes. (F) Number of hydrogen bonds between ApoE and ACE2. (G) Number of hydrophobic interactions between ApoE and ACE2. (H) Interaction energy between ACE2 and ApoE. Brown: ApoE2-ACE2, Green: ApoE3-ACE2 and Blue: ApoE4-ACE2.

**Fig S3.**
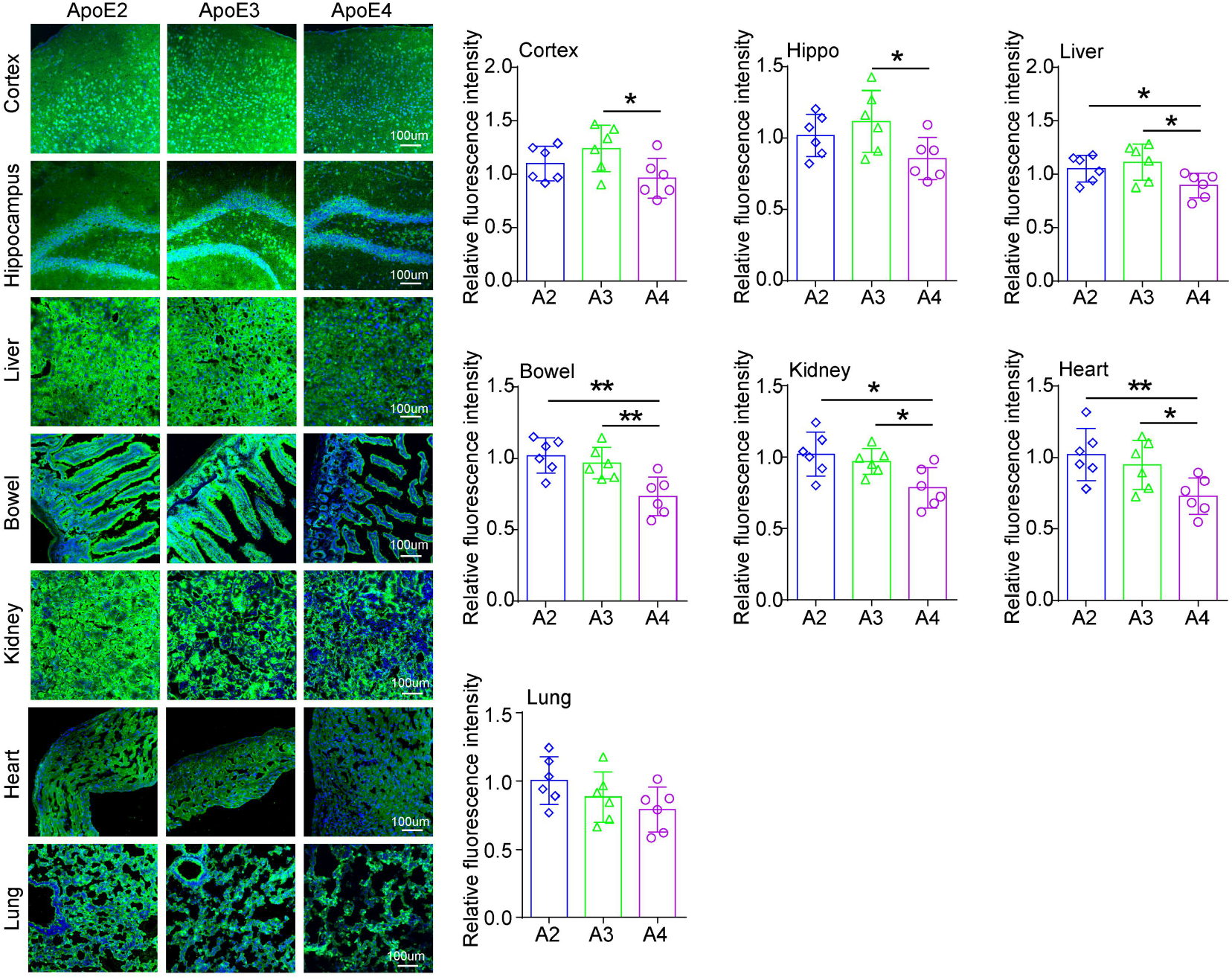
ApoE4 downregulates ACE2 protein expression *in vivo*. ACE2 protein levels in the cortex, hippocampus, liver, bowel, spleen, kidney, heart and lung of ApoE2-TR, ApoE3-TR, and ApoE4-TR mice were assessed by immunofluorescence staining. The results were normalized to the expression of a-tubulin. n = 6 mice per group. The sections were stained with an anti-ACE2 (green) antibody and counterstained with DAPI (blue). The data are expressed as the mean ± SD. Statistical differences were evaluated by one-way ANOVA. Scale bars, 100 μm. \**p* < 0.05; \*\**p* < 0.01; \*\*\**p* <0.001.

**Fig S4.**
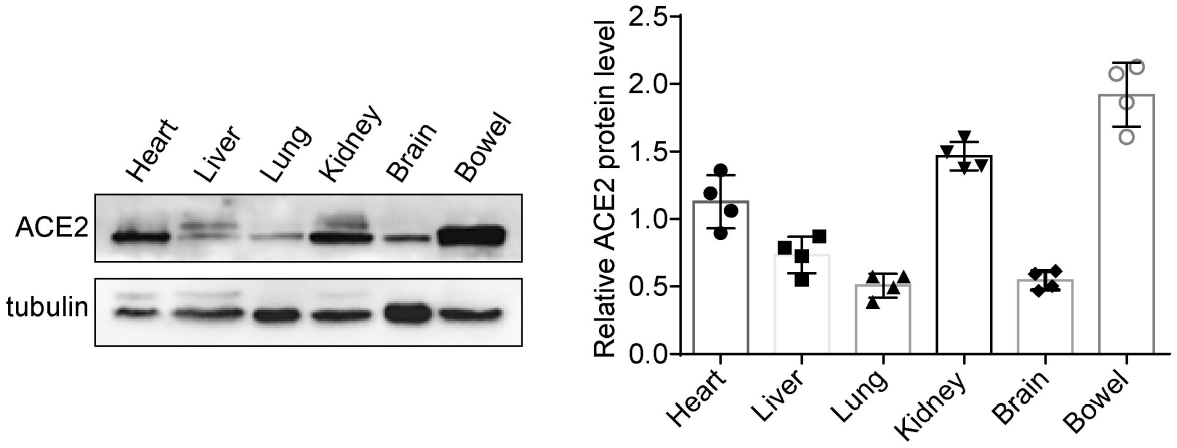
Expression of ACE2 *in vivo*. ACE2 protein levels in the heart, liver, lung, kidney, brain, and bowel (equivalent amounts of total proteins) of ApoE3-TR mice were measured by western blotting, n = 4 mice per group.

**Fig S5.**
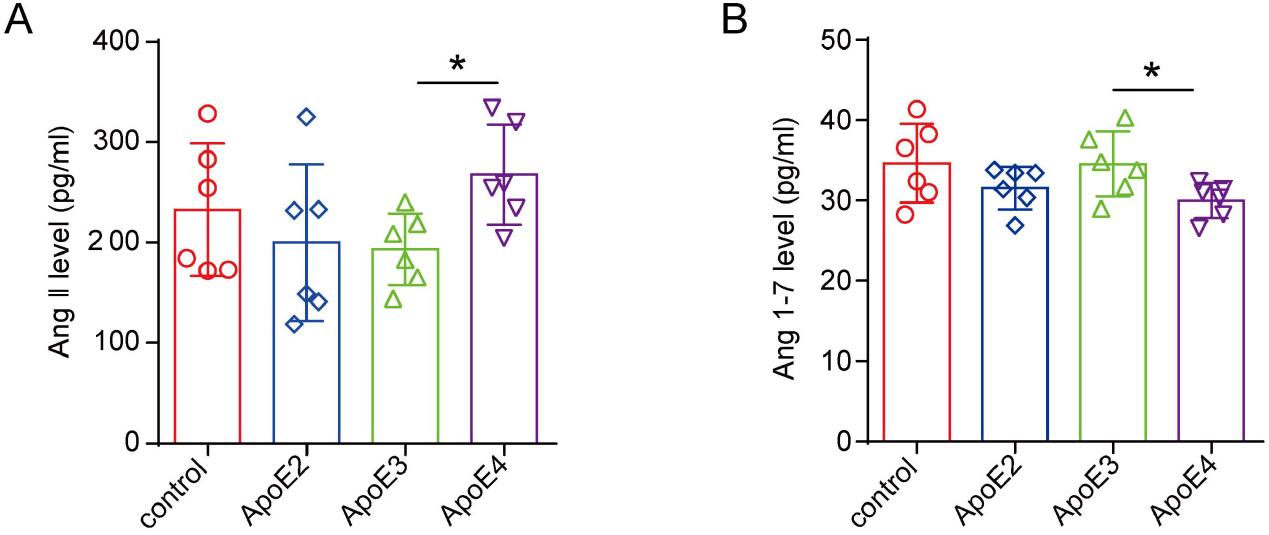
ApoE4 regulates Ang II and Ang 1-7 protein expression in vitro. Expression of the Ang II (A) and Ang 1-7 (B) proteins in HEK-293T cells as shown by ELISA after transfection with 1 µg/ml Flag, ApoE2-Flag, ApoE3-Flag or ApoE4-Flag plasmids for 48 h. The data are expressed as the mean ± SD. One-way ANOVA tests were used. *p<0.05; **p<0.01; ***p<0.001.

**Table S1.**
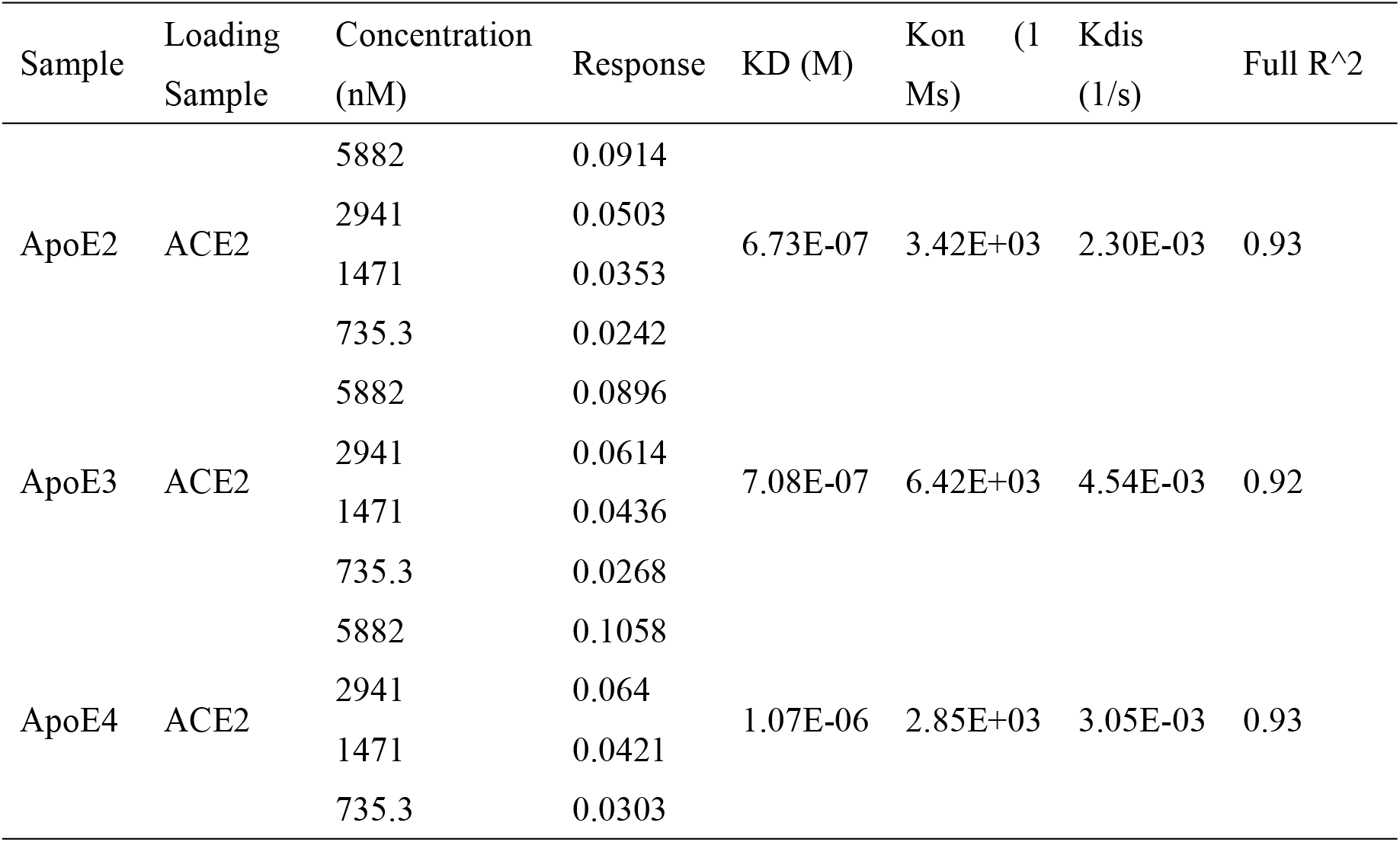
Kinetic parameters of ApoE and ACE2 calculated by BLI.

**Table S2.**
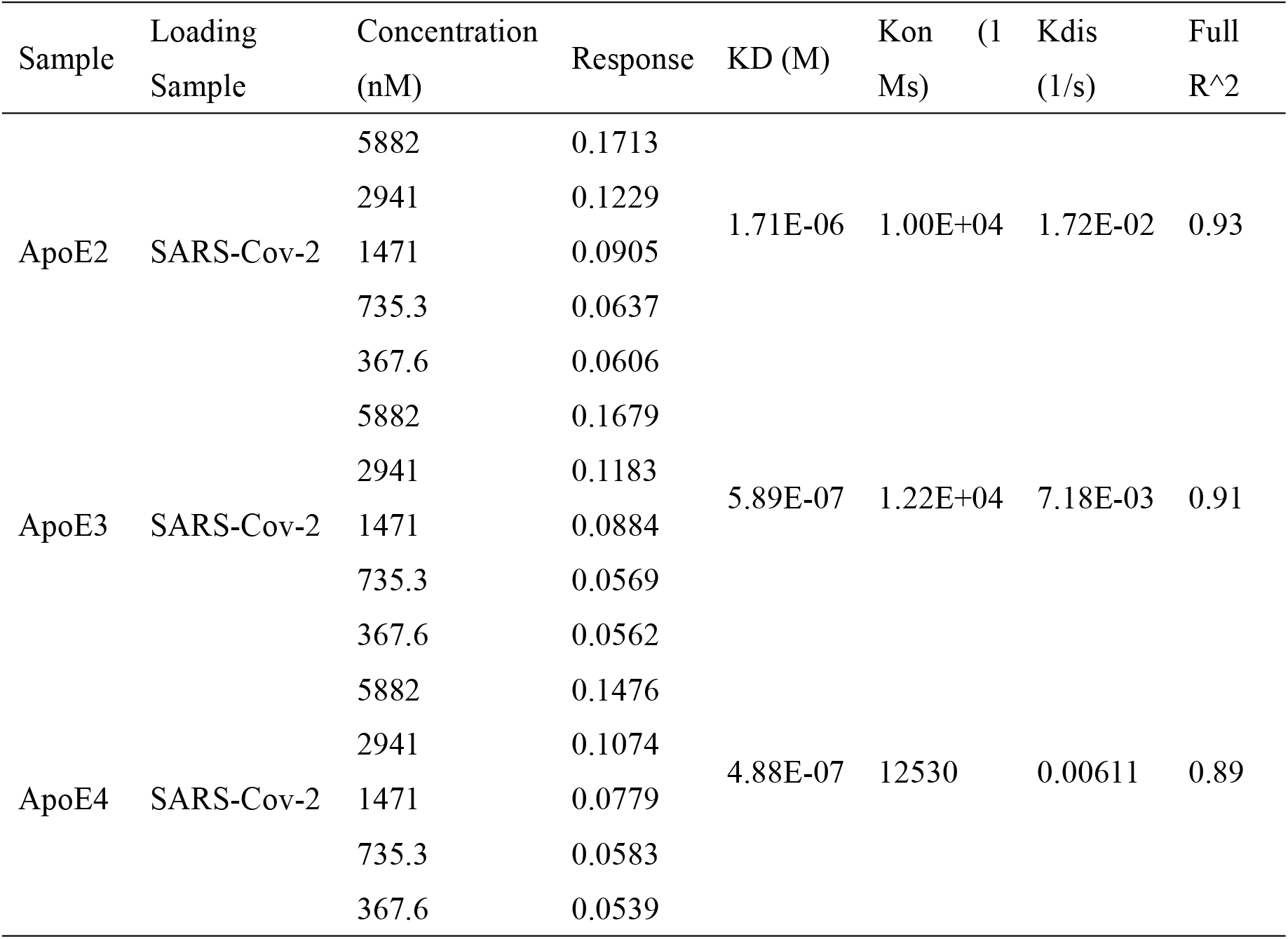
Kinetic parameters of ApoE and SARS-CoV-2 (RBD) calculated by BLI.

## References

1. Clausen TM, et al. (2020) SARS-CoV-2 Infection Depends on Cellular Heparan Sulfate and ACE2. Cell 183(4):1043–1057 e1015.

2. Hamming I, et al. (2004) Tissue distribution of ACE2 protein, the functional receptor for SARS coronavirus. A first step in understanding SARS pathogenesis. J Pathol 203(2):631–637.

3. Hamming I, et al. (2007) The emerging role of ACE2 in physiology and disease. J Pathol 212(1):1–11.

4. Patel VB, Zhong JC, Grant MB, & Oudit GY (2016) Role of the ACE2/Angiotensin 1-7 Axis of the Renin-Angiotensin System in Heart Failure. Circ Res 118(8):1313–1326.

5. Lim S, Bae JH, Kwon HS, & Nauck MA (2021) COVID-19 and diabetes mellitus: from pathophysiology to clinical management. Nat Rev Endocrinol 17(1):11–30.

6. Evans CE, et al. (2020) ACE2 activation protects against cognitive decline and reduces amyloid pathology in the Tg2576 mouse model of Alzheimer’s disease. Acta Neuropathol 139(3):485–502.

7. Wang Y, Li M, Kazis LE, & Xia W (2022) Clinical outcomes of COVID-19 infection among patients with Alzheimer’s disease or mild cognitive impairment. Alzheimers Dement 18(5):911–923.

8. Gordon MN, et al. (2022) Impact of COVID-19 on the Onset and Progression of Alzheimer’s Disease and Related Dementias: A Roadmap for Future Research. Alzheimers Dement 18(5):1038–1046.

9. Severe Covid GG, et al. (2020) Genomewide Association Study of Severe Covid-19 with Respiratory Failure. N Engl J Med 383(16):1522–1534.

10. Kuo CL, et al. (2020) APOE e4 Genotype Predicts Severe COVID-19 in the UK Biobank Community Cohort. J Gerontol A Biol Sci Med Sci 75(11):2231–2232.

11. Kuo CL, et al. (2020) ApoE e4e4 Genotype and Mortality With COVID-19 in UK Biobank. J Gerontol A Biol Sci Med Sci 75(9):1801–1803.

12. Safdari Lord J, Soltani Rezaiezadeh J, Yekaninejad MS, & Izadi P (2022) The association of APOE genotype with COVID-19 disease severity. Sci Rep 12(1):13483.

13. Wang C, et al. (2021) ApoE-Isoform-Dependent SARS-CoV-2 Neurotropism and Cellular Response. Cell Stem Cell 28(2):331–342 e335.

14. Lima FB, Bezerra KC, Nascimento JCR, Meneses GC, & Oria RB (2022) Risk Factors for Severe COVID-19 and Hepatitis C Infections: The Dual Role of Apolipoprotein E4. Front Immunol 13:721793.

15. Kurki SN, et al. (2021) APOE epsilon4 associates with increased risk of severe COVID-19, cerebral microhaemorrhages and post-COVID mental fatigue: a Finnish biobank, autopsy and clinical study. Acta Neuropathol Commun 9(1):199.

16. Zhang H, et al. (2022) APOE interacts with ACE2 inhibiting SARS-CoV-2 cellular entry and inflammation in COVID-19 patients. Signal Transduct Target Ther 7(1):261.

17. Yao X, Gordon EM, Figueroa DM, Barochia AV, & Levine SJ (2016) Emerging Roles of Apolipoprotein E and Apolipoprotein A-I in the Pathogenesis and Treatment of Lung Disease. Am J Respir Cell Mol Biol 55(2):159–169.

18. Martinez-Martinez AB, et al. (2020) Beyond the CNS: The many peripheral roles of APOE. Neurobiol Dis 138:104809.

19. Chen Y, Strickland MR, Soranno A, & Holtzman DM (2021) Apolipoprotein E: Structural Insights and Links to Alzheimer Disease Pathogenesis. Neuron 109(2):205–221.

20. Pu JL, et al. (2022) Apolipoprotein E Genotype Contributes to Motor Progression in Parkinson’s Disease. Mov Disord 37(1):196–200.

21. Zhao J, et al. (2021) Apolipoprotein E regulates lipid metabolism and alpha-synuclein pathology in human iPSC-derived cerebral organoids. Acta Neuropathol 142(5):807–825.

22. Shih IF, Paul K, Haan M, Yu Y, & Ritz B (2018) Physical activity modifies the influence of apolipoprotein E epsilon4 allele and type 2 diabetes on dementia and cognitive impairment among older Mexican Americans. Alzheimers Dement 14(1):1–9.

23. Juul Rasmussen I, Rasmussen KL, Nordestgaard BG, Tybjaerg-Hansen A, & Frikke-Schmidt R (2020) Impact of cardiovascular risk factors and genetics on 10-year absolute risk of dementia: risk charts for targeted prevention. Eur Heart J 41(41):4024–4033.

24. Merello M, Bhatia KP, & Obeso JA (2021) SARS-CoV-2 and the risk of Parkinson’s disease: facts and fantasy. Lancet Neurol 20(2):94–95.

25. Singh AK & Khunti K (2022) COVID-19 and Diabetes. Annu Rev Med 73:129–147.

26. Burt TD, et al. (2008) Apolipoprotein (apo) E4 enhances HIV-1 cell entry in vitro, and the APOE epsilon4/epsilon4 genotype accelerates HIV disease progression. Proc Natl Acad Sci U S A 105(25):8718–8723.

27. Wozniak MA, et al. (2002) Apolipoprotein E-epsilon 4 protects against severe liver disease caused by hepatitis C virus. Hepatology 36(2):456–463.

28. Qiao L & Luo GG (2019) Human apolipoprotein E promotes hepatitis B virus infection and production. PLoS Pathog 15(8):e1007874.

29. Liu S, et al. (2012) Human apolipoprotein E peptides inhibit hepatitis C virus entry by blocking virus binding. Hepatology 56(2):484–491.

30. Kuba K, et al. (2005) A crucial role of angiotensin converting enzyme 2 (ACE2) in SARS coronavirus-induced lung injury. Nat Med 11(8):875–879.

31. Lei Y, et al. (2021) SARS-CoV-2 Spike Protein Impairs Endothelial Function via Downregulation of ACE 2. Circ Res 128(9):1323–1326.

32. Chen J, et al. (2020) Individual variation of the SARS-CoV-2 receptor ACE2 gene expression and regulation. Aging Cell 19(7).

33. Jackson CB, Farzan M, Chen B, & Choe H (2022) Mechanisms of SARS-CoV-2 entry into cells. Nat Rev Mol Cell Biol 23(1):3–20.

34. Schutz D, et al. (2020) Peptide and peptide-based inhibitors of SARS-CoV-2 entry. Adv Drug Deliv Rev 167:47–65.

35. Sivaraman H, Er SY, Choong YK, Gavor E, & Sivaraman J (2021) Structural Basis of SARS-CoV-2-and SARS-CoV-Receptor Binding and Small-Molecule Blockers as Potential Therapeutics. Annu Rev Pharmacol Toxicol 61:465–493.

36. Bu G (2009) Apolipoprotein E and its receptors in Alzheimer’s disease: pathways, pathogenesis and therapy. Nat Rev Neurosci 10(5):333–344.

37. Turner AJ, Hiscox JA, & Hooper NM (2004) ACE2: from vasopeptidase to SARS virus receptor. Trends Pharmacol Sci 25(6):291–294.

38. Monteiro-Junior RS (2020) COVID-19: Thinking about further mental and neurological disorders. Med Hypotheses 143:109894.

39. Gkouskou K, et al. (2021) COVID-19 enters the expanding network of apolipoprotein E4-related pathologies. Redox Biol 41:101938.

40. Yin YW, Sheng YJ, Wang M, Ma YQ, & Ding HM (2021) Interaction of serum proteins with SARS-CoV-2 RBD. Nanoscale 13(30):12865–12873.

41. Camargo SM, et al. (2009) Tissue-specific amino acid transporter partners ACE2 and collectrin differentially interact with hartnup mutations. Gastroenterology 136(3):872–882.

42. Kehoe PG, Wong S, Al Mulhim N, Palmer LE, & Miners JS (2016) Angiotensin-converting enzyme 2 is reduced in Alzheimer’s disease in association with increasing amyloid-beta and tau pathology. Alzheimers Res Ther 8(1):50.

43. Santos RAS, et al. (2018) The ACE2/Angiotensin-(1-7)/MAS Axis of the Renin-Angiotensin System: Focus on Angiotensin-(1-7). Physiol Rev 98(1):505–553.

44. Imai Y, et al. (2005) Angiotensin-converting enzyme 2 protects from severe acute lung failure. Nature 436(7047):112–116.

45. Verdecchia P, Cavallini C, Spanevello A, & Angeli F (2020) The pivotal link between ACE2 deficiency and SARS-CoV-2 infection. Eur J Intern Med 76:14–20.

46. Samavati L & Uhal BD (2020) ACE2, Much More Than Just a Receptor for SARS-COV-2. Front Cell Infect Microbiol 10:317.

47. Riddell DR, et al. (2008) Impact of apolipoprotein E (ApoE) polymorphism on brain ApoE levels. J Neurosci 28(45):11445–11453.

48. Raffai RL, Dong LM, Farese RV, Jr., & Weisgraber KH (2001) Introduction of human apolipoprotein E4 “domain interaction” into mouse apolipoprotein E. Proc Natl Acad Sci U S A 98(20):11587–11591.

49. Montagne A, et al. (2020) APOE4 leads to blood-brain barrier dysfunction predicting cognitive decline. Nature 581(7806):71–76.

50. Shi Y & Holtzman DM (2018) Interplay between innate immunity and Alzheimer disease: APOE and TREM2 in the spotlight. Nat Rev Immunol 18(12):759–772.

51. Zhou Y, et al. (2021) Network medicine links SARS-CoV-2/COVID-19 infection to brain microvascular injury and neuroinflammation in dementia-like cognitive impairment. Alzheimers Res Ther 13(1):110.

